# Quantitative Thermodynamic Characterization of Self-Assembling RNA Nanostructures

**DOI:** 10.1101/2025.07.02.662786

**Authors:** Jordan Aposhian, Surya Pratap S. Deopa, Scott Horowitz, Joseph D. Yesselman

**Author notes:** Correspondence to be addressed to Joseph D. Yesselman and Scott Horowitz.

## Abstract

Recent developments in RNA nanotechnology have led to the rise in designing specific higher order RNA structures with functional goals in mind, such as drug delivery and immunomodulation. As researchers create RNA nanostructures with the goal of becoming common in the molecular biologists’ toolkit, more investigation is required on the robustness of RNA designs. Primarily, what different molecular contexts are the designed and intended nanostructures stable in? In this work we show that by using second-order right-angle light scattering, that RNA nanostructure self-assembly is highly sensitive to environmental conditions. While a test RNA hexagonal grid nanostructure that forms correctly though 120° kissing loops under ideal conditions, small variations in salt conditions and annealing times cause the nanostructure to form less structured variants. Tertiary contacts for self-assembly require magnesium and break over a broad range of low temperatures, melting at 42 °C. In contrast this was found to be considerably lower than the secondary structure melting which occurred at 75 °C. This work underscores the importance of quantitative and thermodynamic characterization of self-assembling nanostructures as they begin to be deployed for engineering and therapeutic applications.x1

## Introduction

Structured RNAs play critical roles throughout the cell by folding into complex three-dimensional (3D) structures that enable a diverse set of functions [1,2]. Building on this natural foundation, researchers have increasingly utilized structured RNAs as blueprints for designing a wide range of nanostructures capable of self-assembly [3–5]. While the original nucleic acid origami systems were DNA-based [6], RNA offers distinct advantages, including co-transcriptional folding [7] that allows these structures to assemble directly within cells [8]. Proposed applications include: RNA nanostructures can deliver siRNA by incorporating targeting elements into their design [9–12], recruit other biomolecules [13], and leverage RNA’s inherent catalytic and functional diversity [14,15]. RNAs ability to form condensates is of great interest to molecular biologists for their ability to concentrate biological reagents [16–18], regulate translation during stress [18–21], and chaperone proteins [22]. Self-assembling nanostructures are also shown to form programmable condensates on their own [23,24].

Despite the growing adoption of self-assembling RNA nanostructures, thermodynamic measurements describing their formation are rare. As these systems become increasingly functional and widespread in molecular biology applications, detailed quantitative characterization of their assembly behavior becomes critical. Currently, the literature provides few methods such as circular dichroism (CD) or native polyacrylamide gel electrophoresis (PAGE) to predict or understand the stability of nanostructure assemblies across varying environmental conditions [25,26]. This knowledge gap will increasingly limit the rate of progress as RNA nanostructures advance toward therapeutic applications, where precise control over assembly and stability is essential. Without such fundamental understanding, the full potential of RNA nanotechnology cannot be realized.

In this study, we develop a comprehensive strategy to measure the thermodynamic stability of self-assembling RNA nanostructures using second-order right-angle light scattering (SORALS) in conjunction with complementary biophysical techniques. Using a model hexagonal grid system (2H_AO_SC) that assembles through 120° kissing loop interactions, we demonstrate that RNA nanostructure assemblies exhibit two distinct melting transitions: secondary structure elements remain stable up to 75 °C. In contrast, tertiary contacts responsible for nanostructure assembly are significantly less stable, melting at 43 °C. We found that these nanostructures require millimolar concentrations of divalent cations and are highly sensitive to solution conditions, forming regular honeycomb patterns only under optimized ionic strength and annealing protocols. However, under non-standardized conditions, the structures form heterogeneous assemblies that maintain tertiary contacts but lack geometric regularity. These findings establish quantitative methods for characterizing RNA nanostructure stability and reveal the critical importance of environmental conditions in controlling assembly behavior, providing insights for the rational design and therapeutic application of RNA nanotechnology.

## Materials and Methods

### DNA Generation

The DNA template of each RNA was primer assembled via PCR obtained through IDT (100 mM in IDTE Buffer, pH 8.0) (Supplemental Table 1). The templates were generated via primer assembly. Middle oligos (denoted p2 and p3 in each oligo pool), were diluted to 1 µM in RNAse-free water. In 19 µL of RNase-free water, 2 µL of middle primers, p1, and p4 primers were diluted. To this Platinum™ Hot Start PCR Master Mix (13000012) was added to begin PCR, following manufacturer’s instructions an initial denaturation of 94°C for 2 minutes. This was followed by 30 cycles of denaturation at 94 °C (30 s), annealing at 60 °C (30 s), and extension at 72°C (30s) on an Applied Biosystems SimpliAmp Thermal Cycler. Major product band was visualized for the correct size on an Invitrogen E-gel system (G8100) with a 2% Agarose E-Gel (G401002) ran for 10 min, and imaged on a Biorad ChemiDoc Imager System (12003153). DNA was purified using the Zymo DNA Clean and Concentrator-5 kit (D4013). DNA concentration was determined by the A260 signal on a ThermoScientific NanoDrop One spectrophotometer (DN-ONE) and extinction coefficient of 1656200.00 M^-1^cm^-1^.

### In vitro transcription

All DNA templates were transcribed to RNA via in vitro transcription. 25 µL of purified DNA template was combined with final concentration transcription buffer (1X Triton, Spermine), 2.5 mM DTT (Sigma Aldrich), 4 mM NTPs (APEX Bio), 20 mM MgCl_2_ (Sigma Aldrich), 200 Units of T7 Polymerase (New England Biolabs) at a final reaction volume of 100 µL. The transcription solution was incubated at 36 °C for 6 hours to maximize RNA yield. The reaction was purified with RNA Clean & Concentrator 5 (Zymo). The correct RNA product was visualized by diluting to 0.5 µM with RNase-free water and 2X RNA Gel Loading Dye (Thermo Fisher Scientific) on a 4% agarose gel, run at 70 V for 2 hours. RNA concentration was determined by the A260 signal and Beer-Lambert Law on a Thermo Scientific NanoDrop One spectrophotometer (DN-ONE) (Supplemental Figure 1).

### Nondenaturing Gel Electrophoresis

All RNA samples, at a final concentration of 0.5 µM, were heated to 90 °C for 5 minutes and then cooled to 4 °C for 4 minutes to denature them for proper folding. Samples were combined with a final concentration of 5 mM magnesium chloride and allowed to fold for 30 min. A Novex 6% TBE Gel (Themo Fisher Scientific) was pre-ran for 1 hr at 250V before loading samples. Samples were combined with BlueJuice (Thermo Fisher Scientific) loading dye diluted to a final concentration of 1X and was ran for 1 hr. Sample was dyed for 10 min with 50 mL of TBE with 2x SYBR Gold (Thermo Fischer Scientific) to visualize the bands. Gel was imaged on a Biorad ChemiDoc Imager System (12003153), (Supplemental Figure 2).

### Second Order Right Angle Light Scattering

RNA was annealed by heating to 90°C for 4 minutes and then cooled to 4°C for 4 minutes using an Applied Biosystems SimpliAmp Thermal Cycler. RNA was then diluted to a final concentration of 5 µM into 10 mM sodium phosphate buffer (pH 7.5, VWR, currently Avantor) in a Hellma microcuvette (115-F-10-40). A second order light scattering signal was read between 500-700 nm from a 310 nm excitation wavelength. The signal was collected using a Cary Eclipse Fluorescence Spectrophotometer at a medium scan speed and a 5 nm slit size. The temperature was manually controlled with single cell peltier attachment. Magnesium titration experiments were carried out by measuring the solution without any magnesium added, followed by titration with 5 µL of magnesium chloride (stocks of 41.5, 81, 83, 160, 342, 693, and 1400 mM), and then waiting 15 minutes for the solution to fold. Thermal measurements were performed by scanning emission from 500-700 nm while exciting at 310 nm. The temperature was then changed and allowed to equilibrate for 1 minute after reaching the desired temperature before the next measurement was taken.

### Atomic Force Microscopy

10 µL of RNA was annealed by heating to 90°C for 4 minutes and cooled to 4°C for 4 minutes in an Applied Biosystems SimpliAmp Thermal Cycler. While heating, AFM Buffer (Tris-Borate 1X (pH 8.1), 2 mM Mg(OAc)_2_, 50 mM KCl, 50 mM NaCl) (Sigma Aldrich) was placed on a freshly cleaved mica surface and preheated to 45 °C. RNA was diluted into this AFM Buffer(Figure 1a, 1b, Supplemental Figure 3, 4, 5, 6, 7) and allowed to cool to 30 °C over the course of 90, 60, or 30 min. RNA was imaged immediately after in tapping mode using a Cypher ES Atomic Force Microscope with an Olympus Biolever Mini (r_nom_ = 8 nm; k_typ_ = 90 pN/nm) at a 25 kHz resonance in liquid. An alternative to this AFM Buffer used was a deposition buffer with the following contents: 10 mM HEPES (pH 7.5), 25 mM KCl, and 10 mM MgCl_2_ (Figure 1c, 1d). The nanostructures in all conditions are of 2nm average height (Supplemental Figures 8, 9), and the large white inclusions are approximately 4 nm high (Supplemental Figure 10). The inclusions are reduced but not eliminated when changing salt conditions and filtering the buffer additional times.

**Figure 1.**
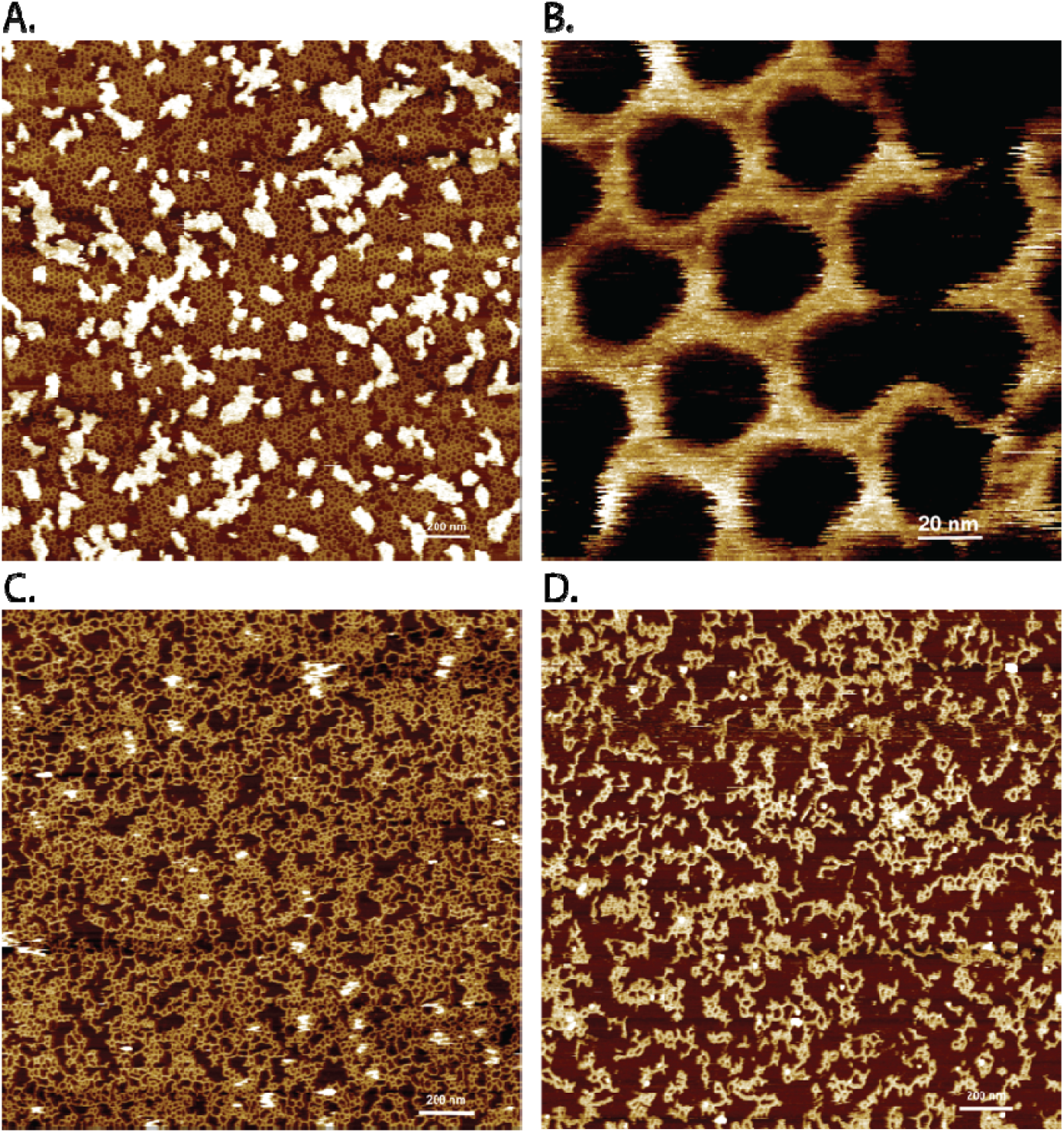
AFM of 2H_AO_SC supporting the fulfillment of tertiary contacts. A) 2 µm x 2 µm image of 250nM annealed for 1hr and 30min in Tris-Borate 1X (pH 8.1), 2 mM Mg(OAc)_2_, 50 mM KCl, 50 mM NaCl showing regular patterned hexagonal formation between tertiary contacts. B) 145nm x 145nm zoom in of same puck. C) 250nM annealed for 1 hr and 30 min in 10 mM HEPES pH 7.5, 25 mM KCl, 10 mM MgCl_2_ shows irregular formation of nanostructures, but tertiary contacts remain fulfilled D) 100nM for 1 hr and 30 minutes.

### Fluorescence Spectroscopy

RNA was heated to 90 °C for 4 minutes and cooled to 4 °C for 4 minutes in an Applied Biosystems SimpliAmp Thermal Cycler. While heating, a 2 mL solution of 10 mM NaPi (pH 7.5) and 1 mM MgCl_2_ was prepared in a Fireflysci (1FLUV10) cuvette. An amount of RNA sufficient for a final concentration of 1 µM was placed in the cuvette and allowed to fold for 30 minutes. After folding, SYBR gold (10,000X in DMSO, Invitrogen) was diluted in the cuvette to a final concentration of 5X. Fluorescence was read every 5 °C in a Cary Eclipse Fluorescence with a medium scan speed and 5 nm slit size spectrophotometer using the built in Agilent Scan Protocol with temperature was controlled manually with single cell peltier attachment.

### Confocal Microscopy

RNA was annealed and heated to 90 °C for 4 minutes, then cooled to 4 °C for 4 minutes, using an Applied Biosystems SimpliAmp Thermal Cycler. RNA was diluted in 10 mM MgCl_2_ and 150 mM HEPES (Sigma Aldrich) and allowed to fold for 30 min. 10 µL of 25 µM folded RNA was placed onto a Cellvis Glass Bottom Dish (D35-10-1.5-N). RNA was imaged under the brightfield to look for coacervation. 1uL of 10X SYBR Gold (Thermo Fisher Scientific) was pipetted into the RNA droplet and visualized using the Alexa Fluor 488 setting at 60x magnification.

### Curve Analysis

Thermodynamic analysis of magnesium titration and melt curve data was performed with a custom Python script, which can be found at https://github.com/JayAposhian/2025NanoPaper. The 4-variable hill equation analysis was performed with GraphPad Prism 6.

## Results

### Model RNA self-assembling nanostructure loses homogeneity in non-standardized conditions

To systematically analyze the folding and structural properties of self-assembling RNA nanostructures, we employed 2H_AO_SC as our model system [7]. This nanostructure is composed of 2 horizontally stacked helical regions (2H) connected by “antiparallel odd” crossover motifs (AO), which can self-assemble (SC). They assemble through 120° kissing loop interactions with four neighboring copies to form hexagonal grid structures [27,28]. Previous work demonstrated that 2H_AO_SC can spontaneously assemble into extended sheets containing regular hexagonal openings under optimal conditions (Figure 1a). To verify that 2H_AO_SC forms the intended 120° kissing loop connections, we characterized nanostructure assembly using atomic force microscopy (AFM). Under optimized (AFM Buffer, see methods) 2H_AO_SC assembled into regular honeycomb structures with well-defined hexagonal openings (Figure 1b) and predicted 2 nm helical diameter (Supplemental Figure 8), consistent with previous reports by Geary et al [7].

However, deviating from these optimized conditions by substituting 1X Tris-Borate pH 8.1 for HEPES buffer 10 mM, pH 7.5, reducing the monovalent salt concentrations from 50 mM NaCl and 50 mM KCl to 25 mM KCl, and increasing 2 mM Mg(OAc)_2_ to 10 mM MgCl_2_ disrupted the structural regularity. This salt condition change resulted in heterogeneous assemblies (Figure 1c, Supplemental Figure 9). Similar irregular structures predominated across multiple annealing time points tested (Supplemental Figure 3-5), suggesting that heterogeneous assembly represents the typical outcome under non-optimized solution conditions. These irregular assemblies likely still maintain kissing loop interactions but lack the geometric organization of the designed honeycomb formation (Figure 1d). These results demonstrate that while 2H_AO_SC can achieve its designed regular architecture, successful assembly requires precise optimization of ionic strength, buffer composition, and thermal annealing protocols.

### 2H_AO_SC reveals discrete melting transitions for secondary and tertiary structure

The thermodynamic stability of RNA nanostructure self-assembly remains poorly characterized in most systems. Using 2H_AO_SC, we systematically examined both secondary and tertiary structure stability. To ensure that observed assembly behaviors arise specifically from kissing loop interactions rather than nonspecific aggregation, we designed a negative control construct (2H_AO_uucg) in which the 120° kissing loops were mutated to UUCG tetraloops that cannot form tertiary interactions [29,30]. We first performed fluorescence-based thermal melting experiments using SYBR Gold. This intercalating dye exhibits enhanced fluorescence only when bound between stacked nucleotide bases, allowing us to monitor the disruption of double-helical secondary structures at elevated temperatures. Both 2H_AO_SC and the 2H_AO_uucg control exhibited steep fluorescence decreases at approximately 75°C (Figure 2a), indicating similar secondary structure melting temperatures regardless of kissing loop functionality.

**Figure 2.**
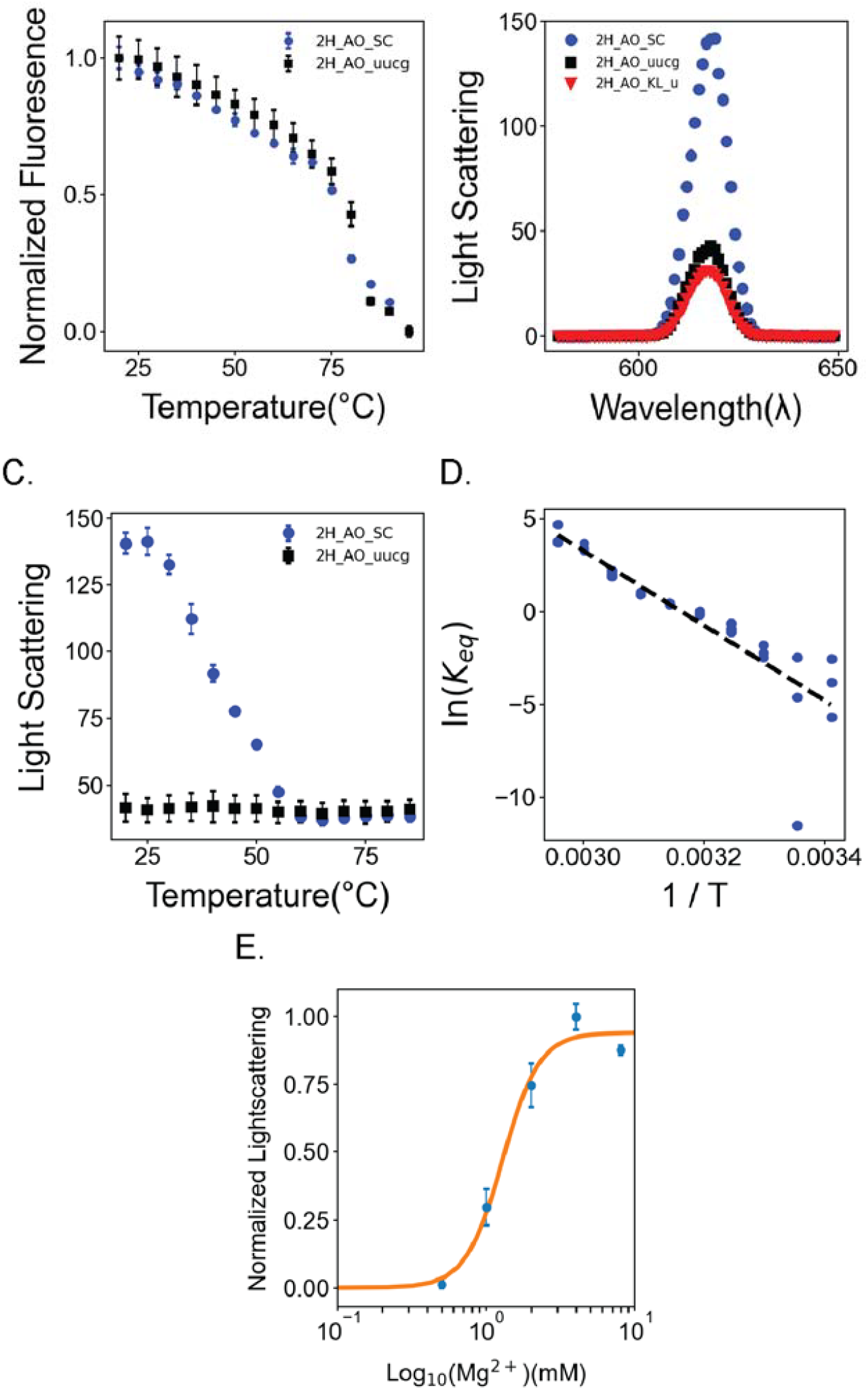
Melting data supports 2 separate transitions for the melting of fully assembled 2H_AO_SC. A) Fluorescence melt over 20°C to 95°C of 1µM 2H_AO_SC and 2H_AO_uucg where both structures melt around 80°C. B) Second order light scattering signal of 5 µM 2H_AO_SC, 2H_AO_uucg, and 2H_AO_KL_at wavelength 620nm, where 2H_AO_SC has a3 times larger signal, indicating a larger particle size. C) Second order light scattering melt over 20°C to 90°C of 5 µM 2H_AO_SC and 2H_AO_uucg where the melt for 2H_AO_SC occurred at 43°C and no significant light scattering signal change was observed for 2H_AO_uucg D) Second order melt temperature data plotted as a Vant Hoft graph E) Second order light scattering signal at 620nm when titrating magnesium to total concentration of 0, 0.5, 1, 2, 4, 8, 16, and 32mM MgCl_2_ into 5 µM 2H_AO_SC, data is fit to a 3 variable hill equation.

However, no differences were observed between the melts of 2H_AO_SC and 2H_AO_uucg, indicating that the fluorescence-based approach lacks the sensitivity required to detect the melting of intermolecular tertiary contacts responsible for nanostructure assembly. This necessitates the use of alternative biophysical methods for characterizing the stability of kissing loops.

To determine the stability of tertiary contacts, we employed second-order right-angle light scattering (SORALS), which provides a sensitive measure of relative particle size and can detect changes in nanostructure assembly [31,32]. To validate SORALS as a method for distinguishing assembled from unassembled nanostructures, we compared the light scattering signals of three constructs: 2H_AO_SC, 2H_AO_uucg, and 2H_AO_KLu. The 2H_AO_KLu construct contained mutations in the middle base pair of all 7-nucleotide kissing loops, disrupting tertiary contact formation. Under folding conditions, 2H_AO_SC exhibited a 3-fold larger scattering signal than both control constructs (Figure 2b), confirming that increased particle size correlates with successful self-assembly.

SORALS analysis revealed a single two-state melting transition for 2H_AO_SC with a T_m_ of 43 °C, approximately 30 °C lower than the secondary structure melting temperature (Figure 2c). This value is consistent with the melting temperature of previously measured small kissing loops [33,34]. As expected, the 2H_AO_uucg control construct exhibited minimal light scattering signal equivalent to the fully dissociated state of 2H_AO_SC across all temperatures (Figure 2C), consistent with the observed signal reports on tertiary contact formation.

These results demonstrate that intermolecular tertiary contacts are significantly less thermostable than intramolecular secondary structures in 2H_AO_SC. The melting profiles also reveal distinct cooperative behaviors: tertiary structure dissociation occurs gradually over approximately 30 °C, indicating a heterogeneous population of assembly states, while secondary structure melting proceeds cooperatively over a narrow 5-10 °C range. Van’t Hoff analysis of the tertiary structure melting transition yielded thermodynamic parameters of ΔH = -149 kJ/mol and ΔS = 0.5 kJ/mol·K (Figure 2d, see methods Curve Analysis). The self-assembly is confirmed to be an exothermic process, where heat is released as noncovalent interactions form between the nanostructure monomers.

### 2H_AO_SC requires divalent ions to self-assemble

Another important characteristic of nanostructures is that they require ions due to the high charge density in folded RNAs, which is necessary for RNA to achieve teritary structure [35–37]. It was unclear how much would be needed to form this grid-type nanostructure. We investigated the dependence of these nanostructures on cation concentration.

Light scattering intensity was measured as a function of magnesium concentration, revealing a standard saturation binding curve where magnesium rapidly increased scattering from 0 to 400 intensity units. 2H_AO_SC achieved its maximum assembly size at 1.5 mM Mg^2+^, while the 2H_AO_uucg control reached only 29% of this maximum signal when titrated with 1.5 mM Mg (Supplemental Figure 11, Supplemental Table 2). From this data, we calculated an Mg1/2 of 1.28 mM Mg2+, consistent with literature values for magnesium requirements of folded RNAs (Supplemental Figure 1, Supplemental Table 2) [38]. Hill equation analysis yielded a Hill coefficient of 1, indicating non-cooperative binding (Figure 2e). These experiments demonstrate that light scattering provides a valuable tool for characterizing RNA nanostructure stability under varying ionic conditions.

### 2H_AO_SC forms disordered condensates

Recent studies have demonstrated that self-assembling nucleic acid nanostructures can undergo liquid-liquid phase separation (LLPS) [39–41]. Furthermore, nucleic acids are the core of condensates in cells [42] as well as being an essential part of dictating LLPS properties [43]. Nucleic acid-protein LLPS have also been implicated in many diseases, such as neurodegenerative diseases [44–46]. This prompted us to investigate whether our model self-assembling RNA nanostructures could form condensates. Confocal microscopy visualization of 2H_AO_SC using SYBR Gold at low magnification revealed no discrete coacervates but instead showed large, disordered structures exceeding 0.1 mm in length. This structural disorder resembles the irregular networks observed in AFM imaging (Figure 3a). Notably, the SYBR signal localized exclusively to the glass surface rather than remaining in solution, indicating that 2H_AO_SC undergoes surface-mediated condensation rather than remaining freely dispersed. In contrast, 2H_AO_uucg exhibited only diffuse SYBR fluorescence under identical conditions (Figure 3b), while SYBR alone showed no signal as expected (Figure 3c). This combination suggests that the condensed 2H_AO_SC is concentrating the SYBR into its condensed form on the glass and depleting the SYBR from any monomer that remains in solution, or that very little monomer remains in solution and that the majority has been incorporated into the large irregular condensate. Combined, these results show that this model RNA nanostructure condenses into an irregular structure that forms sizes close to a macro-scale, approximating the width of a human hair.

**Figure 3.**
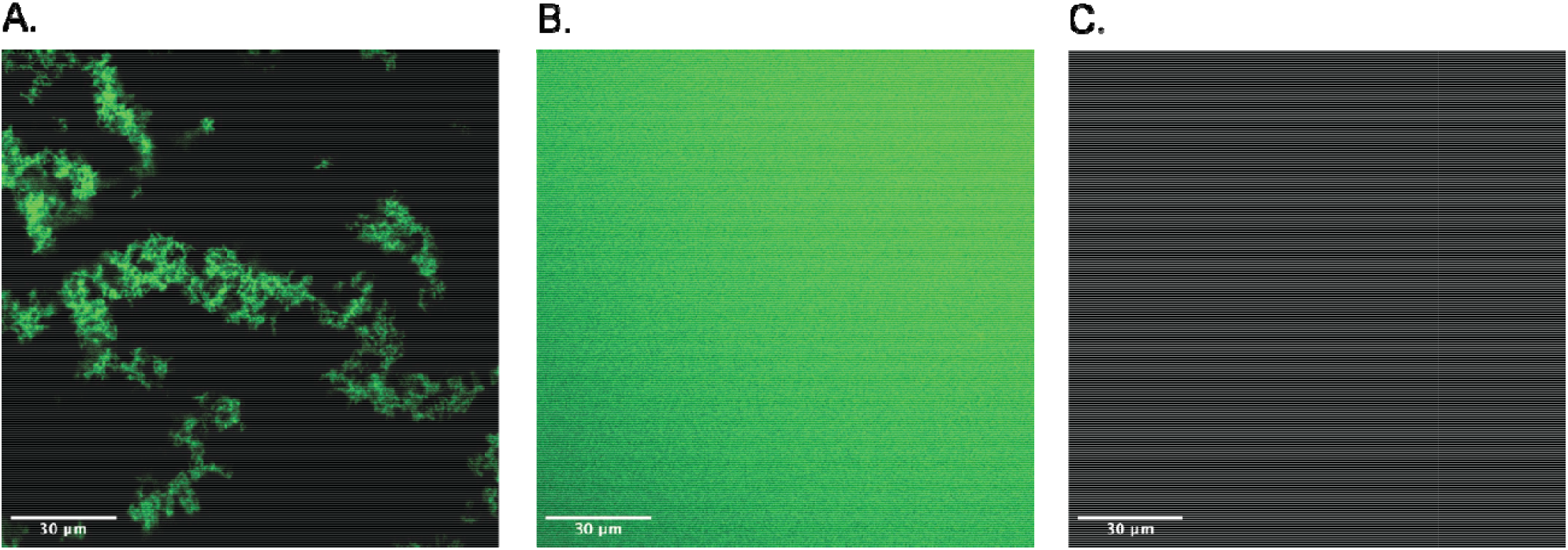
Confocal microscopy imagining reveals structural homology. A) 40 µM 2H_SC_AO in 150 mM HEPES and 10 mM MgCl_2_ with SYBR Gold showing distinct, concentrated areas of SYBR fluorescence B) 40 µM 2H SC_AO in 150 mM HEPES and 10 mM MgCl_2_ with SYBR Gold showing diffuse SYBR signal spread around the slide C) in 15 mM HEPES and 10 mM MgCl_2_ with SYBR Gold shows no signal.

## Discussion

The characterization of RNA nanostructure thermodynamic stability has been limited by the lack of accessible, label-free analytical methods that can distinguish between the contributions of secondary and tertiary structure to assembly formation. As RNA nanostructures advance toward applications in drug delivery, immunology, and synthetic biology [47–50], understanding their stability limitations becomes critical for rational design and predicting performance under diverse experimental conditions. The development of standardized analytical approaches will provide essential tools for researchers implementing these systems in cellular and therapeutic contexts.

Our SORALS-based approach demonstrates the utility of light scattering for quantitative stability analysis under controlled ionic conditions. This approach represents a rapid, label-free analytical method that can directly measure the thermodynamics of tertiary contacts across diverse solution conditions. The observed tertiary contact melting temperature of 43°C, approximately 30°C lower than that of secondary structure elements, is consistent with previous measurements of kissing loops, such as the HIV-1 dimerization initiation sequence (DIS). Importantly, we demonstrated SORALS’ sensitivity to magnesium concentration and buffer composition changes, making it a powerful screening tool for systematically evaluating how varying ionic conditions—including those approaching physiological relevance—affect nanostructure assembly behavior.

A key finding of this work is that RNA nanostructures exhibit significant assembly heterogeneity under non-optimized conditions, forming irregular but functional assemblies rather than the regular geometric patterns observed under carefully controlled protocols. This heterogeneity likely represents the predominant state of these systems in many experimental contexts and suggests that nanostructures rarely achieve the stoichiometric, geometrically perfect assemblies predicted by computational design approaches [51]. Understanding this assembly landscape is crucial for predicting how these systems will behave in complex biological environments where precise control over ionic conditions and annealing protocols is not feasible.

Future development of this methodology should focus on expanding the analytical toolkit to include measurements of chemical stability, nuclease resistance, and behavior under varying solvation conditions—all critical parameters for therapeutic applications. Additionally, increasing the sensitivity of current assays to detect subtle structural perturbations would enable high-throughput analysis of mutational libraries, accelerating the optimization of nanostructure designs for specific applications.

## Acknowledgements

This work was funded by the This work was funded by NIH R35GM142442 (S.H.), This work was supported by the NSF(NSF CAREER 214363) to JD.Y. We would like to thank Tom Perkins Ph.D. for the facilities and equipment to do the Atomic Force Microscopy studies. Graphical abstract created in BioRender. Horowitz, S. (2025) https://BioRender.com/8sqmxn0

## CRediT authorship contributions statement

**Jay Aposhian:** Investigation, Formal Analysis, Visualization, Writing – original draft, Writing – review & editing **Surya Pratap S. Deopa:** Investigation, Visualization, Writing-review & editing **Joseph Yesselman**: Conceptualization, Data Curation, Funding Acquisition, Project Administration, Supervision, and writing – review & editing **Scott Horowitz:** Conceptiontualization, Data Curation, Funding Acquisition, Project Administration, Supervision, and writing – review & editing

