## Supplemental Data for "Quantitative Thermodynamic Characterization of Self-Assembling RNA Nanostructures"

Supplemental Table 1:

| Type | Name | Sequence |
| --- | --- | --- |
| DNA | 2H_AO_SC-p1 | TTCTAATACGACTCACTATAGGAATGAACCTGAAGGAGGCACGGGTACGGGTCGCG |
| DNA | 2H_AO_SC-p2 | GCGGAATGATCAGGCGAGACGTGCCTGAACGGGTCGCACGTCTCGCGACCCGTACC |
| DNA | 2H_AO_SC-p3 | CGCCTGATCATTCCGCACAGGTCGGTGAAGCCTCCACGCCGAGCTCGACCCGTCCACTGG |
| DNA | 2H_AO_SC-p4 | GCACAGGGCCTACGTCCACTGTAGGCGCTCGACCCAGTGGACGGGTCGAGCTCGGCGT |
| DNA | 2H_AO_SC_uucg-p1 | TTCTAATACGACTCACTATAGGAATGAACCTGAAGGAGGCACGGGTACGGGTCGCTT |
| DNA | 2H_AO_SC_uucg-p2 | CGGAATGATCAGGCCGAAGCCTGAACGGGTCGCCGAAGCGACCCGTACCCGTGCC |
| DNA | 2H_AO_SC_uucg-p3 | CGGCCTGATCATTCCGCACAGGTCGGTGAAGCCTCCACGCCGAGCTCGACCCTTCGGGG |
| DNA | 2H_AO_SC_uucg-p4 | GCACAGGGCCTACCGAAGTAGGCGCTCGACCCCGAAGGGTCGAGCTCGGCGTGG |
| DNA | NetMut_p1_JA | TTCTAATACGACTCACTATAGGAATGAACCTGAAGGAGGCACGGGTACGGGTCGCGAGT |
| DNA | NetMut_p2_JA | GCGGAATGATCAGGCGAGACGTGCCTGAACGGGTCGCACGACTCGCGACCCGTACC |
| DNA | NetMut_p3_JA | CGCCTGATCATTCCGCACAGGTCGGTGAAGCCTCCACGCCGAGCTCGACCCGTCTACT |
| DNA | NetMut_p4_JA | GCACAGGGCCTACGTCAACTGTAGGCGCTCGACCCAGTAGACGGGTCGAGCTCGGCGT |
| RNA | 2H_AO_SC | GGAAUGAACCUGAAGGAGGCACGGGUACGGGUCGCGAGACGUGCGACCCGUUCAGGCACGUCUCGCCUGAUCAUUCCGCACAGGUCGGUGAAGCCUCCACGCCGAGCUCGACCCGUCCACUGGGUCGAGCGCCUACAGUGGACGUAGGCCCUGUGC |
| RNA | 2H_AO_SC_uucg | GGAAUGAACCUGAAGGAGGCACGGGUACGGGUCGCUUCGGCGACCCGUUCAGGCUUCGGCCUGAUCAUUCCGCACAGGUCGGUGAAGCCUCCACGCCGAGCUCGACCCUUCGGGGUCGAGCGCCUACUUCGGUAGGCCCUGUGC |
| RNA | NetMut | GGAAUGAACCUGAAGGAGGCACGGGUACGGGUCGCGAGUCGUGCGACCCGUUCAGGCACGUCUCGCCUGAUCAUUCCGCACAGGUCGGUGAAGCCUCCACGCCGAGCUCGACCCGUCUACUGGGUCGAGCGCCUACAGUUGACGUAGGCCCUGUGC |

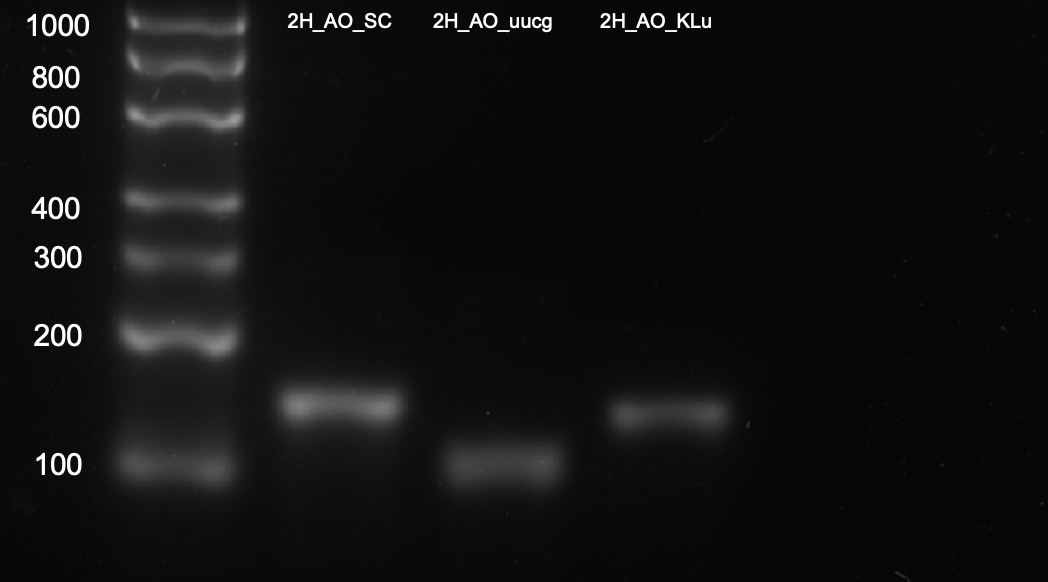

Supplemental Figure 1: Denaturing Agarose Gel Electrophoresis of RNA Nanostructures, 4% agarose gel with formamide, ladder given in single stranded base pairs

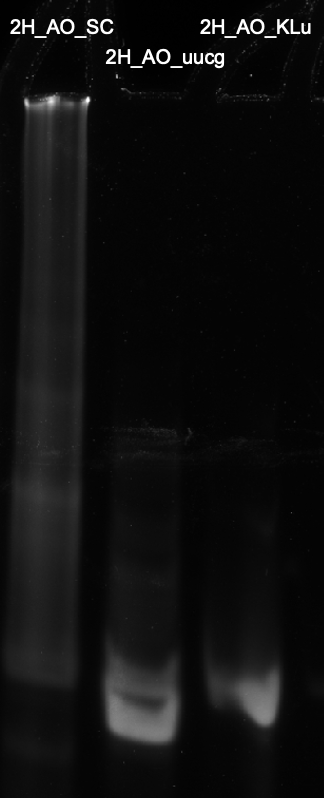

Supplemental Figure 2: Nondenaturing polyacrylamide gel electrophoresis of 1uM 2H_AO_SC, 2H_AO_uucg, 2H_AO_KLu, 6% TBE Gel with 5mM magnesium chloride in folding buffer and in gel buffer.

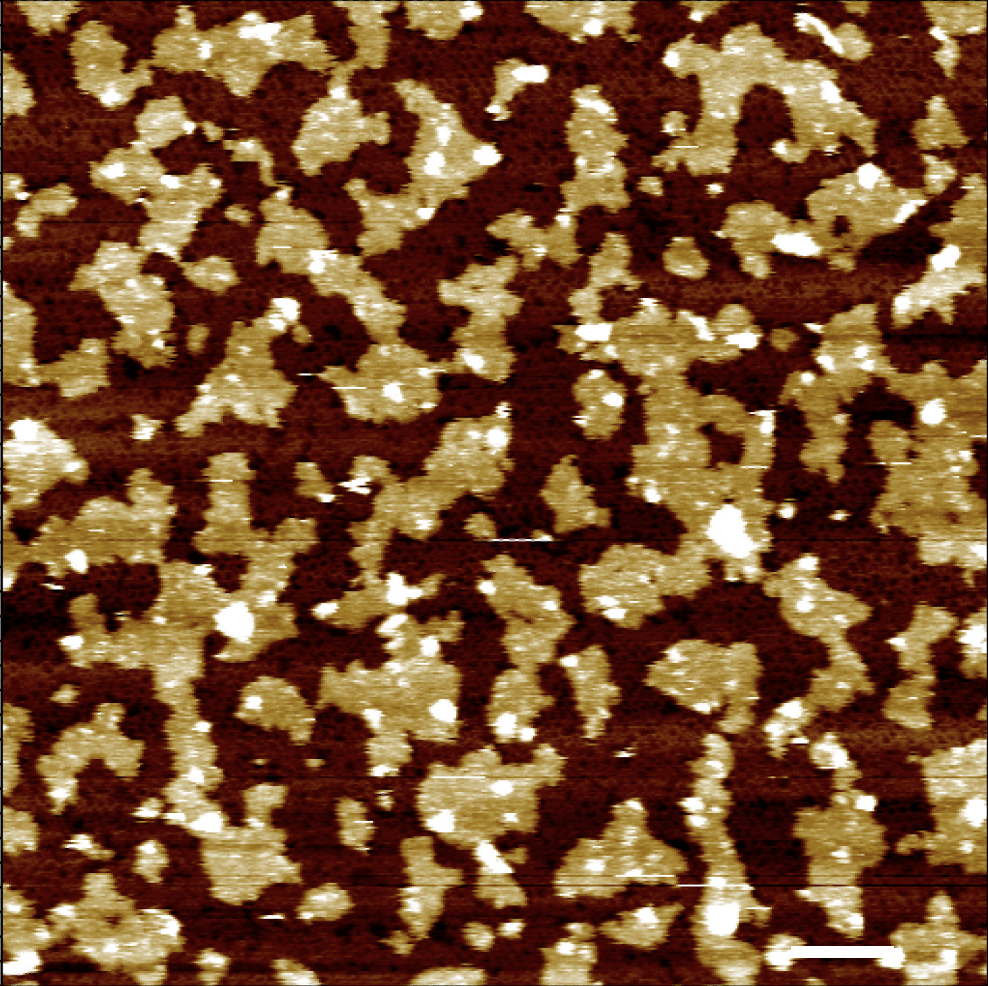

Supplemental Figure 3: Condition: AFM Buffer- 30 minute anneal, Scale Bar 200 nm

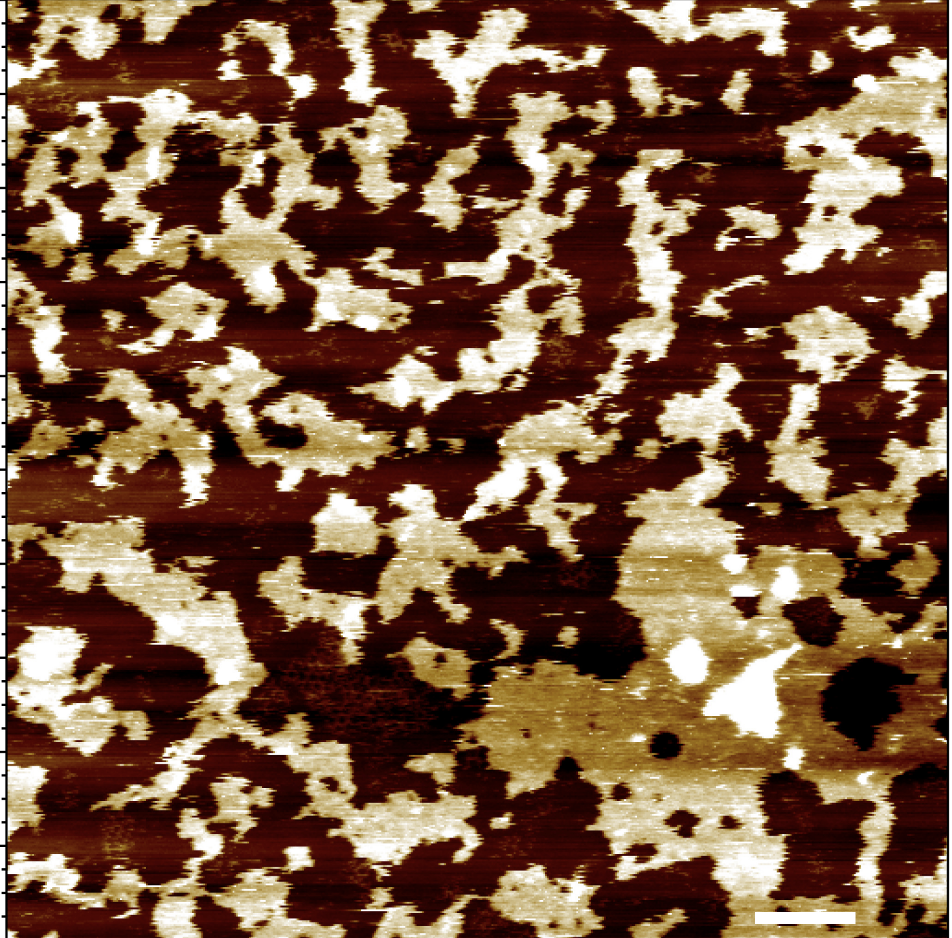

Supplemental Figure 4: Condition AFM Buffer- 30 minute anneal 10 nm, scale bar 200 nm

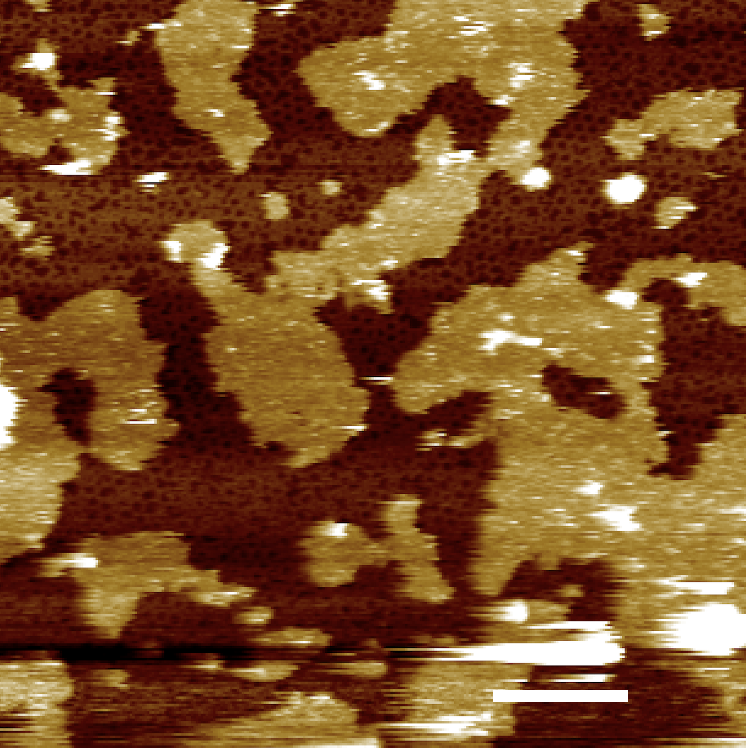

Supplemental Figure 5: Conditions AFM Buffer- 1 hour anneal, 100 nm, scale bar 200 nm

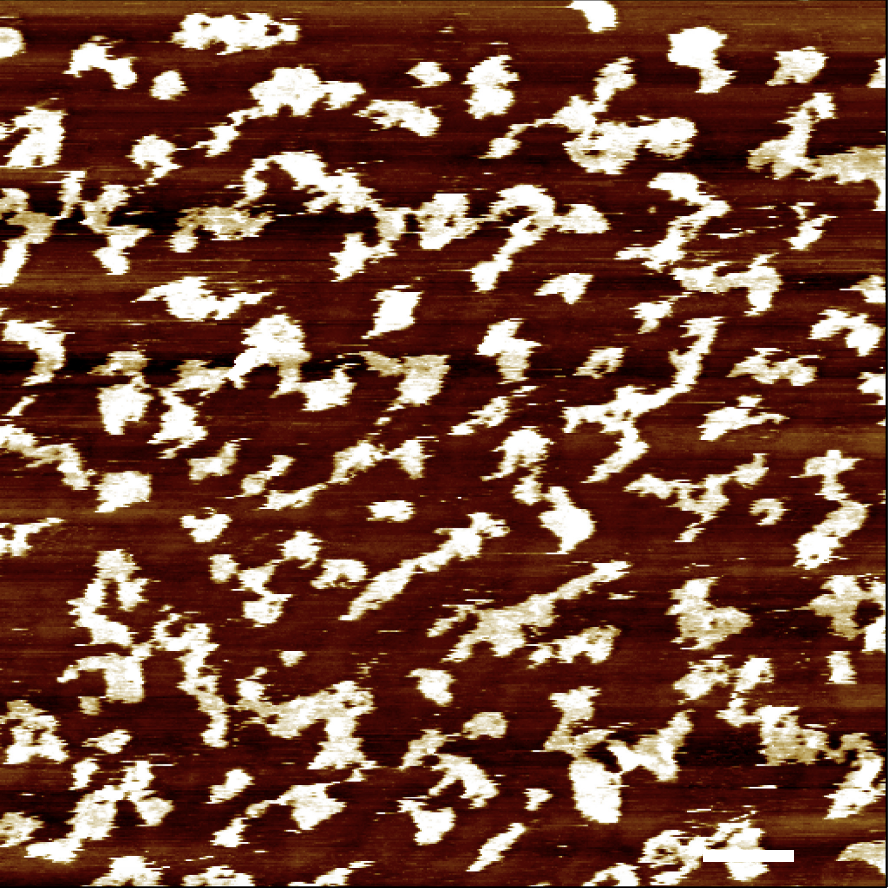

Supplemental Figure 6: Conditions AFM Buffer, 30 minutes anneal, buffer alone, scale bar 200 nm

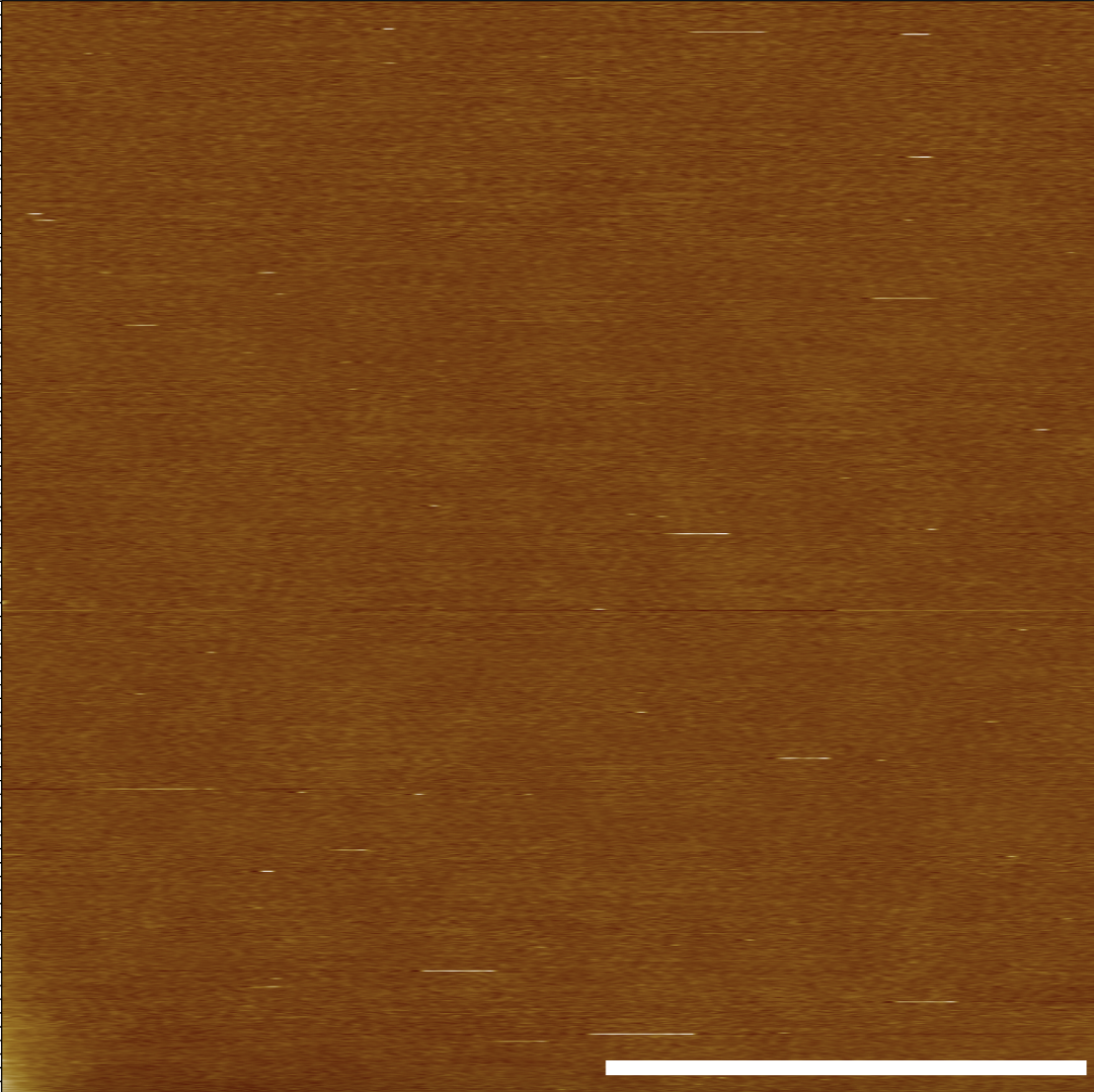

Supplemental Figure 7: Conditions AFM Buffer, 30 minutes anneal, water, scale bar 200 nm

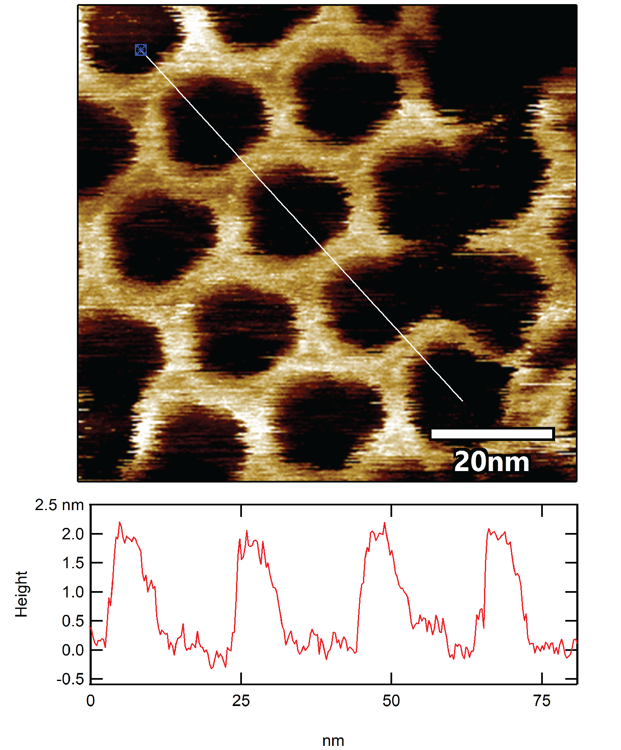

Supplemental Figure 8: Height graph of Figure 2B showing even 2 nm height of 2H_AO_SC on atomic force microscopy

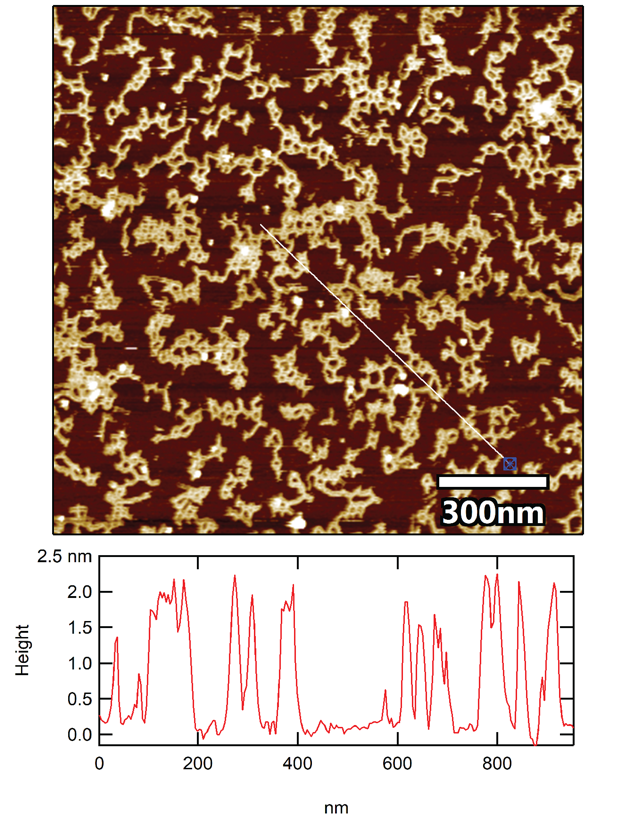

Supplemental Figure 9: Height graph of Figure 1D showing even 2 nm height of 2H_AO_SC on atomic force microscopy

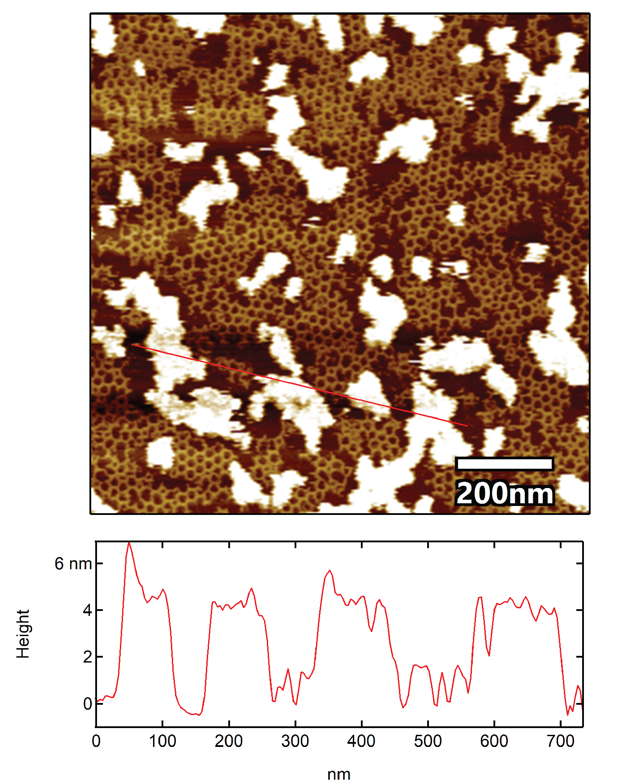

Supplemental Figure 10: Height graph of Figure 1A showing average 4 nm height of inclusions with 2 nm high 2H_AO_SC

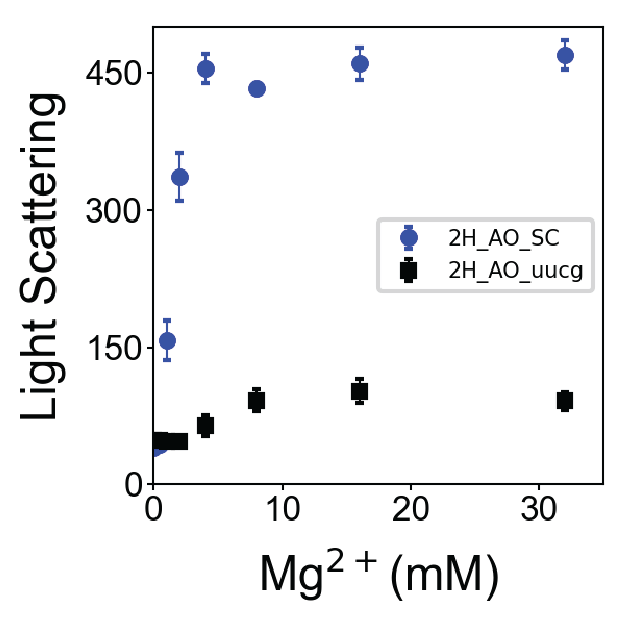

Supplemental Figure 11: Light scattering of magnesium titration of 2H_AO_SC and 2H_AO_uucg

| log(agonist) vs. response -- Variable slope (four parameters) | |
| --- | --- |
| Best-fit values | |
| Bottom | 0 |
| Top | 376 |
| LogEC50 | 1.358 |
| HillSlope | 1.023 |
| EC50 | 22.81 |
| Span | 376 |
| Std. Error |  |
| Top | 10.78 |
| LogEC50 | 0.09678 |
| HillSlope | 0.1862 |
| 95% Confidence Intervals | |
| Top | 348.3 to 403.8 |
| LogEC50 | 1.109 to 1.607 |
| HillSlope | 0.5449 to 1.502 |
| EC50 | 12.86 to 40.45 |
| Goodness of Fit | |
| Degrees of Freedom | 5 |
| R square | 0.99 |
| Absolute Sum of Squares | 2052 |
| Sy.x | 20.26 |
| Constraints |  |
| Bottom | Bottom = 0.0 |
| Number of points | |
| Analyzed | 8 |

Supplemental Table 2: Data and Fit Parameters associated with 4 Variable Hill Equation Fit

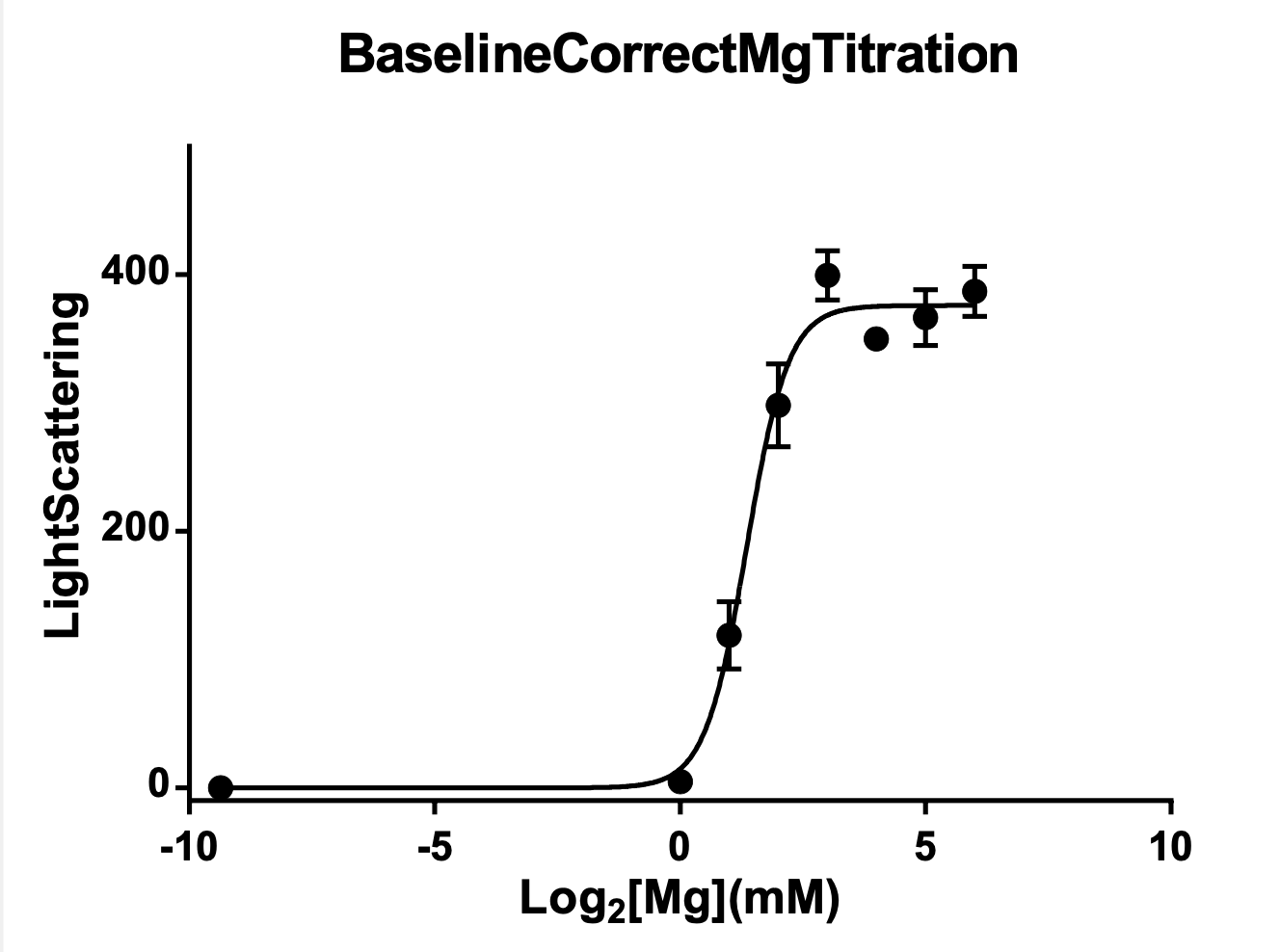

Supplemental Figure 12: 4 Variable Hill Equation Magnesium Titration Fit
